# Genome-wide identification and characterization of the NAC transcription factor family in *Cynodon dactylon* and their expression during abiotic stresses

**DOI:** 10.64898/2026.04.15.718725

**Authors:** Anit Poudel, Yanqi Wu

## Abstract

Common bermudagrass (*Cynodon dactylon*) is a highly resilient and cosmopolitan grass widely used for turf, forage, and soil stabilization. Although its genome has been sequenced, little study has focused on characterizing genes underlying its resilience, including the NAC transcription factor family, which is well known for its physiological and stress-related functions. This study aimed to systematically characterize NAC TF genes in the bermudagrass genome and assess their potential roles in abiotic stress tolerance. A total of 237 CdNAC genes were identified and phylogenetically classified into 14 groups, including 40 members in the NAM/NAC1 class, which is associated with plant growth and development, and 23 members in the SNAC class, which is associated with stress responses. Tissue-specific RNA-seq analysis indicated that about one-fourth of CdNAC genes were expressed across all tissues, whereas 13 genes showed relatively higher expression in roots and 9 in inflorescence, suggesting both essential and specialized functions. Stress-responsive expression profiling revealed that 35 CdNAC genes were upregulated in response to drought, 43 to heat, 10 to salt, and 42 to submergence stress. Notably, CdNAC122, 149, and 155, the members of SNAC class, were consistently upregulated across all stress conditions, while others exhibited stress-specific expression, such as CdNAC37, 130, 145, and 199 in drought, CdNAC7, 12, 18, and 29 in heat, CdNAC46 and 151 in salt, and CdNAC9 and 31 in submergence. In contrast, 53 genes were downregulated during different stresses, with most belonging to NAM/NAC1, TERN, or OsNAC7 classes, possibly reflecting suppression of photosynthesis and development-related processes under stress. These results provide the first comprehensive characterization of CdNAC genes, reveal their distinct regulatory roles in abiotic stress responses, and establish a foundation for future functional validation and applications in breeding of stress-resilient bermudagrass.

## Introduction

Plants grow and reproduce in a complex environment characterized by a wide range of abiotic factors. Climatic and edaphic factors, including temperature, light, water availability, soil composition, and salinity, are critical determinants of plant growth, development, reproduction, and survival. Fluctuations in these factors beyond their optimal ranges can disrupt normal physiological and biochemical processes. For instance, high salt concentration limits water absorption, induces osmotic and oxidative stress, and causes ion toxicity [1]. High temperature disrupts membrane integrity, causes ion leakages, and impairs photosynthesis and respiration [2]. Water deficit reduces water use efficiency, causes cellular dehydration, and impairs normal growth and development [3]. Similarly, submergence stress limits oxygen availability and hinders cellular respiration [4].

Nevertheless, plants have evolved intricate mechanisms to mitigate these stresses, including signal transduction pathways that activate stress-responsive genes. Transcription factors (TFs) play a critical role in this regulatory network by regulating gene expression in response to environmental signals. Among diverse transcription factors found in plants, NAC (No Apical Meristem [NAM], *Arabidopsis* Transcription Activation Factor [ATAF1/2], and Cup-Shaped Cotyledon 2 [CUC2]) is one of the largest families of plant-specific TFs, highly reported for their response to abiotic stresses alongside their critical role in cellular growth and development. Its role has been well documented for developmental functions such as embryogenesis, meristem development, secondary wall formation, xylem differentiation, leaf senescence, root, shoot, floral, fruit, and seed development, as well as stress responses such as drought, salinity, heat, cold, flooding, nutrient deficiency, and pathogen stresses [5].

NAC protein typically consists of a highly conserved N-terminal DNA-binding domain and a variable C-terminal region. The N-terminal domain is structurally organized into five subdomains (A-E) that are critical for DNA binding and dimerization, while the C-terminal region–also known as the transcriptional activation region (TAR)–plays a role in protein interactions and transcriptional activation or repression of target genes. Some NAC proteins also contain atypical α-helical transmembrane domain at the C-terminus, known as NAC with a transmembrane motif1-like (NTL) proteins, which are often anchored to plant membranes and believed to have roles in sensing stress signals [6,7].

*Cynodon dactylon* (common bermudagrass) is a warm-season perennial grass widely grown for turf, forage, and soil conservation due to its rapid growth, extensive root system, dense ground cover, and high traffic tolerance [8,9]. While bermudagrass exhibits notable resilience to various environmental challenges, its overall performance is still significantly constrained by abiotic stresses such as prolonged drought or flooding, high salinity, and extreme temperatures. Recent breeding efforts have focused on enhancing abiotic stress tolerance, mostly through trait-based selection, to enhance quality and diversify uses while conserving water [10,11]. However, to achieve faster and more precise genetic gains, identifying genes involved in stress tolerance has become important.

Though the importance of NAC TFs in stress responses has been established in several plant species, this family remains unexplored in bermudagrass. In this study, we aim to identify and characterize the complete NAC gene family in *C. dactylon*, analyze their gene structures, conserved motifs, and evolutionary relationships, and investigate their expression pattern under various abiotic stresses. We hypothesize that polyploidy-driven expansion of NAC genes contributes to abiotic stress resilience in bermudagrass. This comprehensive understanding of the NAC gene family in *C. dactylon* could provide valuable insights for breeding programs aimed at enhancing stress tolerance, as well as future studies on functional roles and regulatory pathways of NAC TFs in this ecologically and economically important grass species.

## Materials and methods

### Identification and chromosomal mapping of bermudagrass NAC transcription factors

The genomic sequence, General Feature Format (GFF) annotation, coding sequence, and protein sequence of *Cynodon dactylon* (PRJCA019991) were downloaded from the Genome Warehouse (https://ngdc.cncb.ac.cn/gwh/) database. The HMM profile of the NAC domain (PF02365) from the Pfam (http://pfam.xfam.org/) database was used to search for bermudagrass NAC proteins using HMMER v3.4 (http://hmmer.org) with an E-value threshold of 1e^-5^. The redundant sequences were manually removed, and the sequences were further verified for the presence of complete NAC domains using NCBI CDD batch search (https://www.ncbi.nlm.nih.gov/Structure/bwrpsb/bwrpsb.cgi) and SMART (http://smart.embl-heidelberg.de/). The corresponding chromosome position of identified NAC genes was extracted from the GFF file, and their chromosome distribution was visualized using MapGene2Chrom (http://mg2c.iask.in/mg2c_v2.0/).

### Physicochemical properties and subcellular localization

The physical and chemical properties, including molecular weight and isoelectric point of CdNACs were estimated using the Biopython package [12]. Subcellular localization of all CdNAC proteins was estimated using the web-based server Cello (https://cello.life.nctu.edu.tw/). The presence of transmembrane helix in CdNACs was predicted using DeepTMHMM [13].

### Gene structure, conserved motif, and cis-element analysis

The gene structure of CdNACs was analyzed using the Gene Structure Display Server (https://gsds.gao-lab.org/). The conserved motifs in CdNAC proteins were identified using MEME Suite (https://meme-suite.org/meme/) and visualized using TBtools-II v2.331[14].

To predict cis-acting elements of CdNAC genes, 2000 bp sequences upstream of the transcription start site in the promoter region of each CdNAC gene were extracted using TBtools-II and submitted to PlantCare (https://bioinformatics.psb.ugent.be/webtools/plantcare/html/).

### Phylogenetic, Syntenic, and Evolutionary Analysis

The NAC protein sequences of bermudagrass were aligned with *Arabidopsis* NAC protein sequences using Clustal Omega [15], and a phylogenetic tree was constructed using the maximum likelihood method in RAxML [16] with 1000 bootstraps and visualized using itol [17].

The gene duplication events and syntenic relationship of CdNACs were analyzed using the MCScanX [18]. We used pairwise BLASTp at an e-value threshold of 10^-10,^ keeping the top 15 hits to analyze the duplication events. Non-synonymous (Ka) and synonymous (Ks) nucleotide substitution rates for each duplicated pair were estimated using TBtools-II, and their ratio was used to evaluate the selective pressure. The approximate evolutionary time of the duplication event was estimated using T=Ks/2λ, where λ=6.5 x 10^-9^ substitutions per nucleotide per year [19]. We also compared the genome sequence of *C. dactylon* with its grass relatives *Oropetium thomaeum* and *Oryza sativa* to understand the evolutionary nature of CdNAC genes based on gene synteny.

### RNA-seq data and expression analysis

To elucidate the expression of CdNAC genes at different tissue levels (root, shoot, rhizome, stolon, leaf, and inflorescence) and different abiotic stresses (heat, drought, salt, and submergence), publicly available genome-wide transcription data were obtained from the NCBI SRA database (https://www.ncbi.nlm.nih.gov/sra) under the accession numbers PRJNA685207 and PRJNA589598. The stress was induced continuously for 7 days in 3-week-old bermudagrass seedlings by withholding water, treating seedlings with 400mM NaCl, submerging the seedlings under water, and exposing seedlings to 42 °C, thereby extracting RNA from leaf tissue of stress-induced and control samples with two biological replicates [20]. RNA-seq data were mapped to the *C. dactylon* reference genome using HISAT2 [21], and reads were counted using HTSeq-count [22]. Normalization of RNA-seq read count and differential expressions of genes was analyzed using edgeR [23] and visualized as a heatmap of log2-transformed values using TBtools-II.

## Results

### Identification and chromosome distribution of CdNAC TFs

Initially, a total of 244 non-redundant putative NACs were identified using an HMM search. Using CDD and SMART search, 7 proteins with nonsignificant hits and incomplete NAC subdomains were removed to obtain a final total of 237 NACs. The slightly higher number of NACs in bermudagrass, compared to diploid grass species, may be attributed to its tetraploid nature (2n = 4x = 36) and its role in diverse transcriptional regulatory networks.

We mapped 237 NACs to all 18 chromosomes and named them from CdNAC1 to CdNAC237 based on their chromosomal position. Chromosome 3B had the maximum number of CdNACs (25), whereas chromosome 8A had only 1 CdNAC (Figure 1).

**Figure 1:**
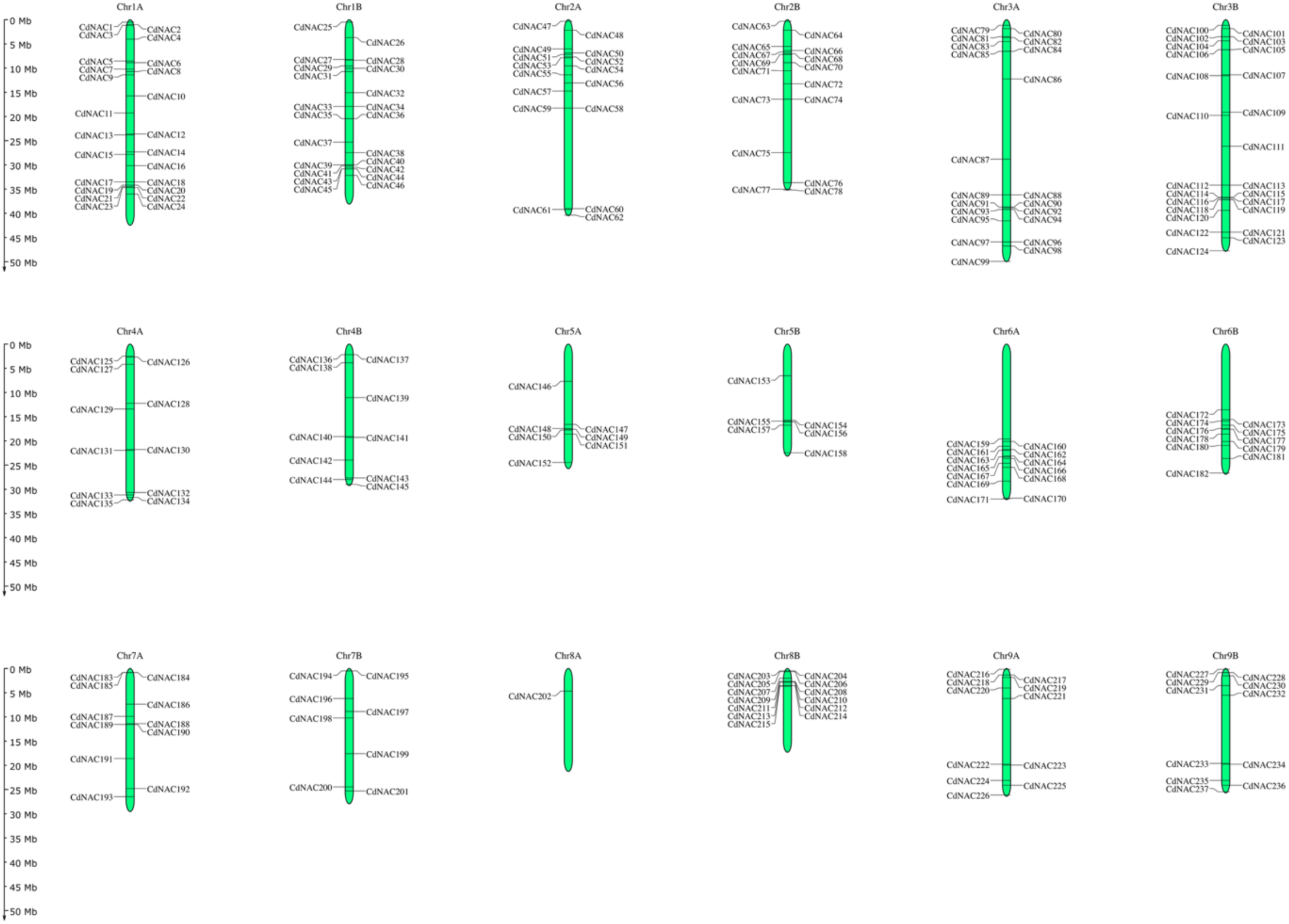
Distribution of CdNACs in 18 chromosomes of common bermudagrass (*Cynodon dactylon*)

The physicochemical properties of CdNAC greatly varied in bermudagrass. The amino acid sequence length of CdNAC ranged from 158 to 904, molecular weight from 18001.0671 to 99232.28 Da, and Isoelectric point from 4.437 to 11.2 (Table S1). Cellular localization analysis estimated that 183 CdNAC proteins were most likely localized in the nucleus, 31 in the cytoplasm, 9 in mitochondria, 8 in the chloroplast, 5 in the extracellular space, and 1 in the vacuole (Table S1). Only two CdNACs, CdNAC53 and CdNAC69, were predicted to contain transmembrane helix at 686-703 and 685-702 amino acid positions, respectively.

### Phylogenetics of CdNAC gene family

The phylogenetic analysis of 237 bermudagrass CdNAC genes, along with 105 *Arabidopsis* ANAC genes, grouped the genes into 14 classes (Figure 2). The grouping of the genes was inspired by Ooka et al. (2004), who classified *Arabidopsis* and rice NAC genes into 18 subclasses based on sequence similarity as well as the differences in motifs present in C-terminal variable regions, which typically depict the functional characteristics of NAC transcription factors. Based on this classification, 23 CdNAC genes were grouped within the SNAC subfamily that contains subgroups like AtNAC3, NAP, and ATAF, which is well described as the stress-responsive NAC gene class in NAC TF analysis [7,24]. Similarly, 40 CdNAC genes were classified in the NAM/NAC1 group, and this group is well studied to have plant development-related functions like shoot apical meristem formation, morphogenesis, and growth [25]. Similarly, four new classes, like ANAC006, ANAC027, CdNAC2, and XND1, were defined based on their high bootstrap support values (Figure S3). CdNAC2 does not host any members from *Arabidopsis* NAC genes, and the XND1 class is believed to impact root hydraulics and trade-off for stress response [26].

**Figure 2:**
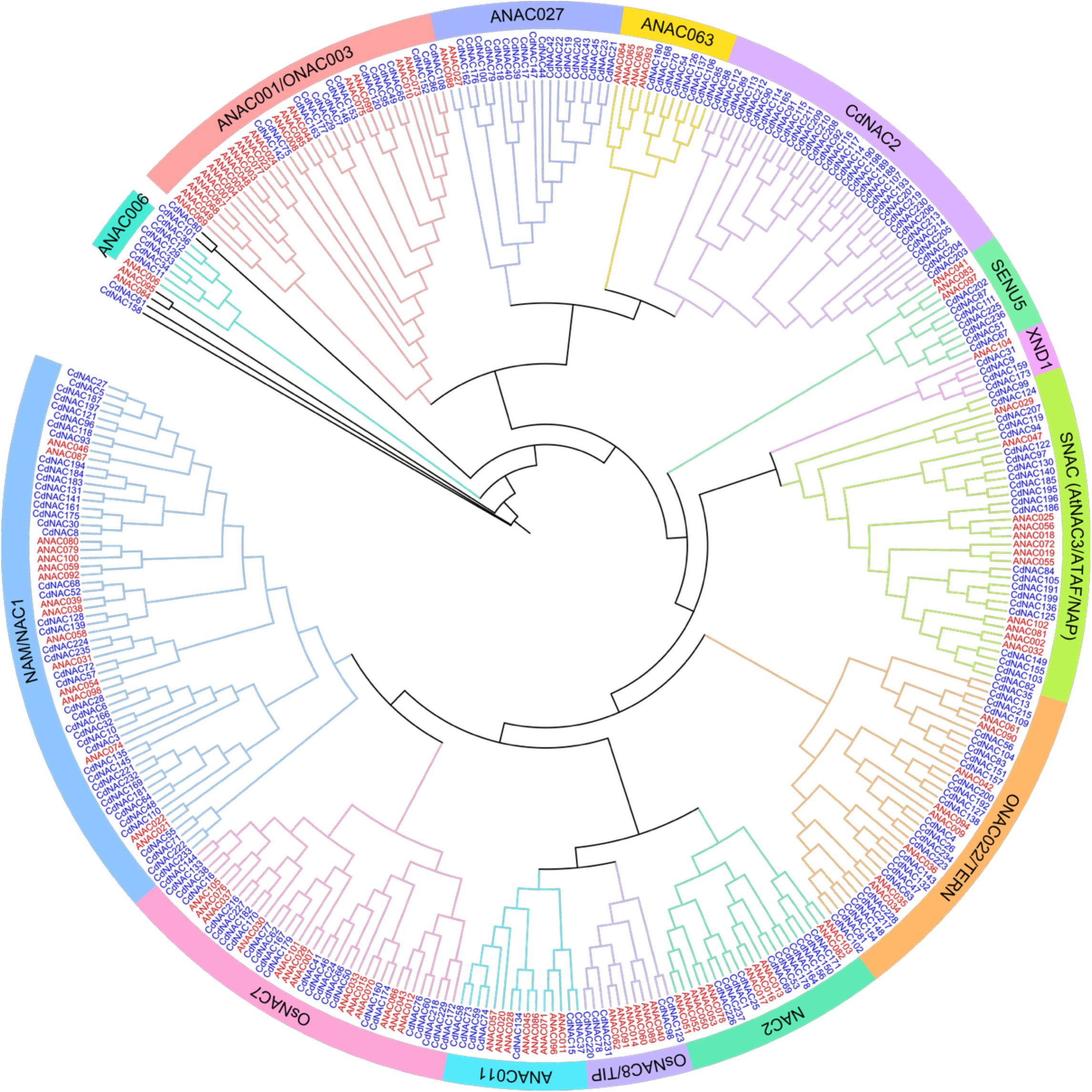
Phylogenetic classification of bermudagrass and *Arabidopsis* NAC proteins

### Gene structure and conserved motifs of CdNAC genes

The study of gene structure and conserved motifs can help elucidate the structural and functional characteristics of CdNAC genes. We used meme software to search for the top 15 conserved motifs. The first 7 motifs were found in most of the CdNAC proteins, and these motifs were located within the five N-terminal sub-domains (A-E) of the NAC domain (Figure S1 and S2). Some motifs were specific to a particular group, supporting the phylogenetic classification. For example, motifs 8 and 10 were specific to ONAC003, motif 14 was more specific to OsNAC7, and motifs 11, 12, and 15 were specific to ANAC027. The results are on par with Ooka et al. (2003) those who have identified 13 common motifs in TARs corresponding to different subgroups. The conserved motif analysis also largely supports the two broad classifications proposed by Ooka et al. (2003), with group II notably lacking motifs 1 and 3. Similarly, members within a phylogenetic class had similar exon numbers and patterns (Figure 3). For instance, members of ANAC027 and CdNAC2 had very few exons, ranging from 1 to 3. Members in XND1 had 3 exons, members of the NAC2 class had 4-5 exons. While members of ANAC001/ONAC003 are among the NAC proteins with the highest exon numbers, ranging from 5-6 to 12.

**Figure 3:**
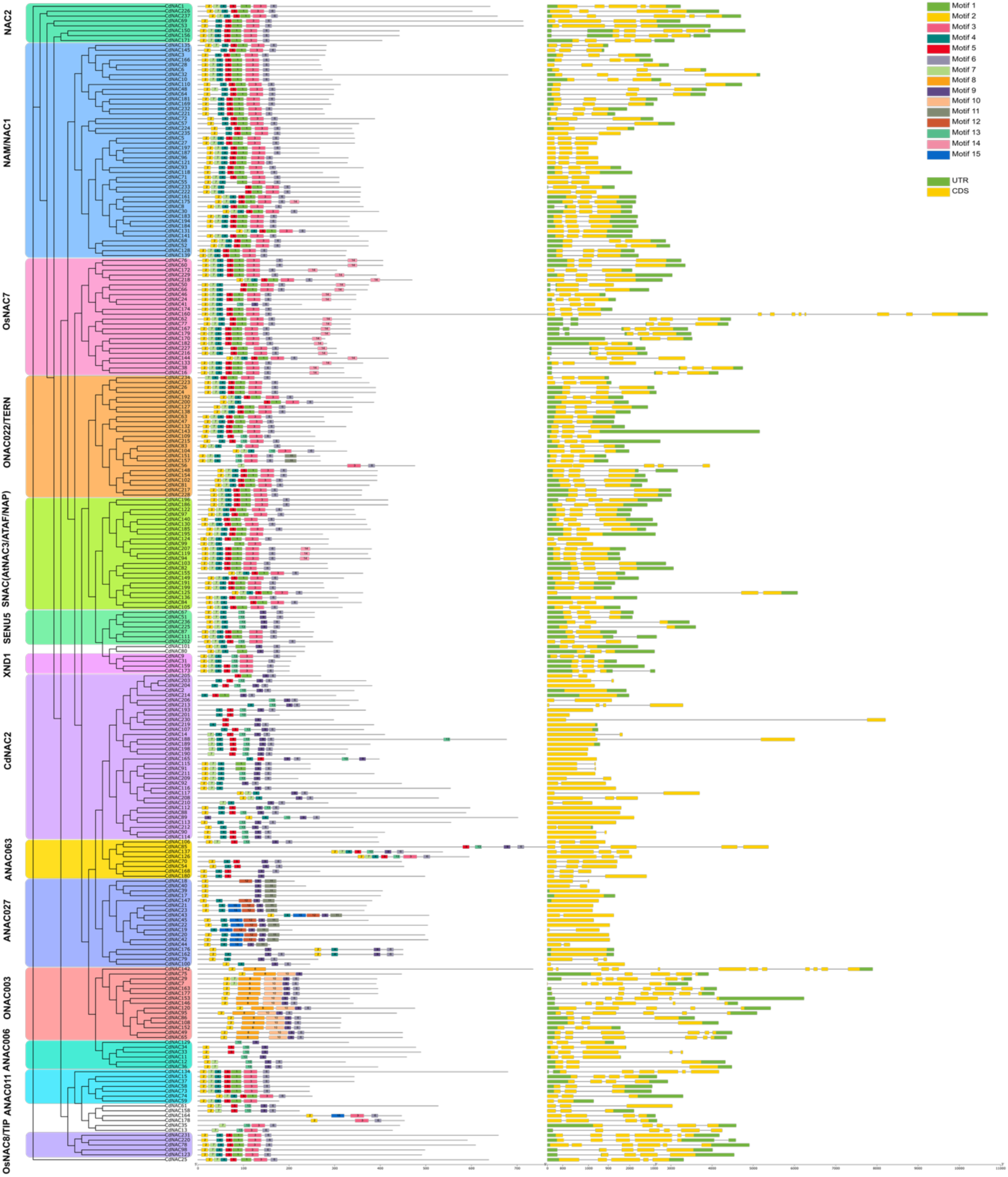
Phylogenetic tree, conserved motif, and gene structure of CdNAC TFs.

### Syntenic and evolutionary analysis of CdNAC genes

To understand the evolutionary relation and expansion of CdNAC family genes, a genome-wide synteny analysis was performed. 109 gene pairs were collinear, involving 147 genes arising from the whole genome or segmental duplication (Table S2). Additionally, 35 CdNAC genes were in tandem duplication, contributing to 38 tandemly duplicated gene pairs. Most of the collinear gene pairs were from homoeologous chromosomes, supporting the tetraploid nature of common bermudagrass with two homologous subgenomes (Figure 4).

**Figure 4:**
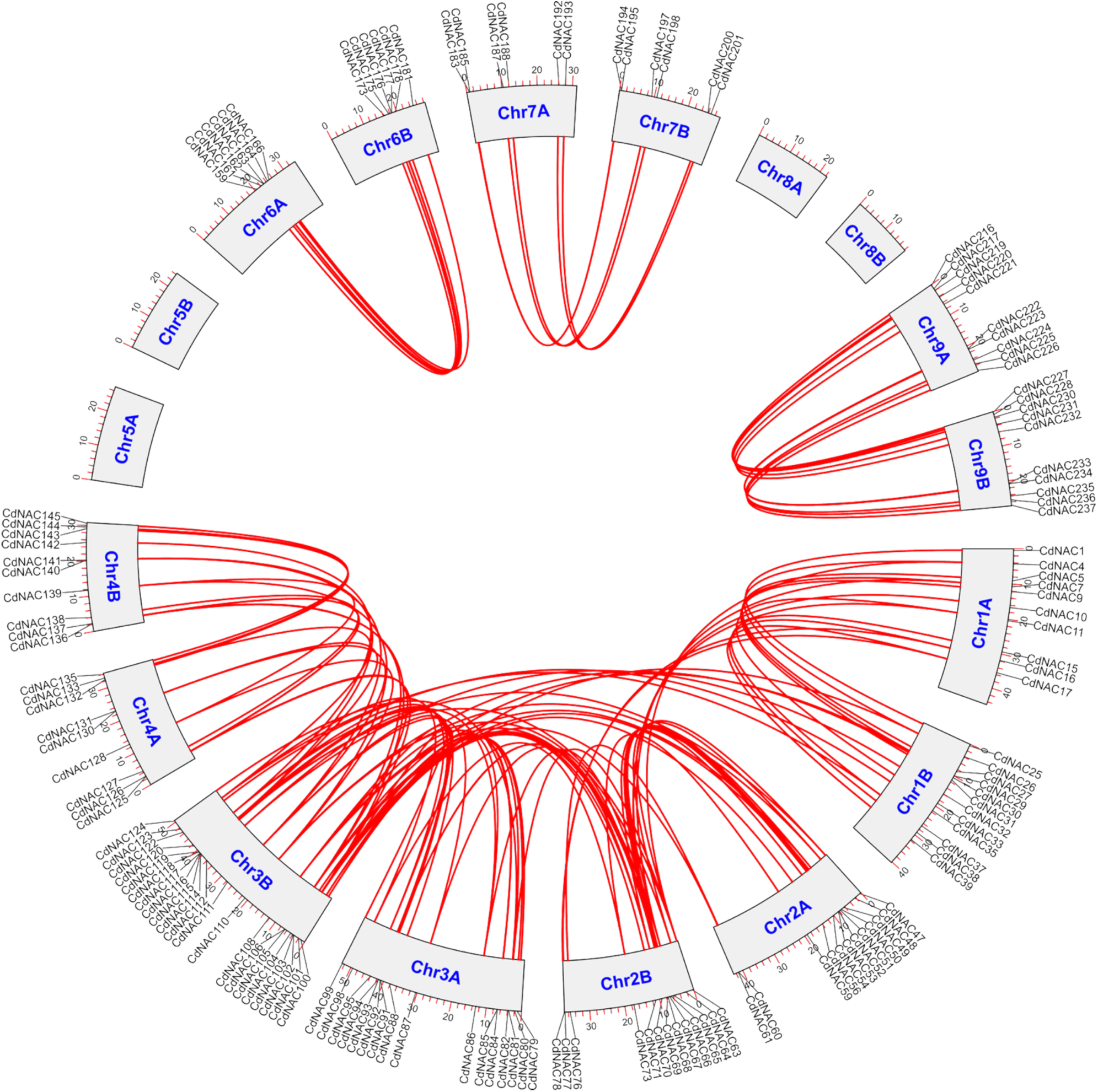
Syntenic analysis of CdNAC genes

The rate of nonsynonymous and synonymous substitution rates reflects the selective pressures on the evolution of duplicated genes. The majority of duplicated gene pairs have a Ka/Ks value less than 1, suggesting purifying (negative) selection between duplicated genes with conserved functional characteristics (Table S2). While 2 gene pairs exhibited a Ka/Ks ratio greater than 1, indicating potential positive selection, suggesting neofunctionalization or adaptive evolution. The divergence time of these duplicated genes is estimated to be between 205.80 million years ago to 0.89 million years ago.

1. T. Fang et al. (2020) utilized single-nucleotide variants to study the genomic evolution of *C. dactylon* and revealed the allotetraploid nature of the plant with a high degree of orthology with Oropetium thomaeum, a highly drought-tolerant resurrection plant commonly used for desiccation studies. Given the strong resilience characteristics of *C. dactylon* and its evolutionary linkage with *O. thomaeum*, we conducted an interspecific collinearity analysis between CdNAC genes and the *O. thomaeum* genome. We identified 240 syntenic gene pairs involving 169 CdNAC genes and 91 *O. thomaeum* genes with a high level of chromosomal correspondence, potentially explaining the expansion of CdNAC genes and retention of functional characteristics. Similarly, we found substantial synteny with *Oryza sativa* genes, where 160 CdNAC genes were syntenic with 95 rice genes, giving 241 collinear pairs, indicating a high degree of conservation of NAC genes in the Poaceae family (Figure 5).

**Figure 5:**
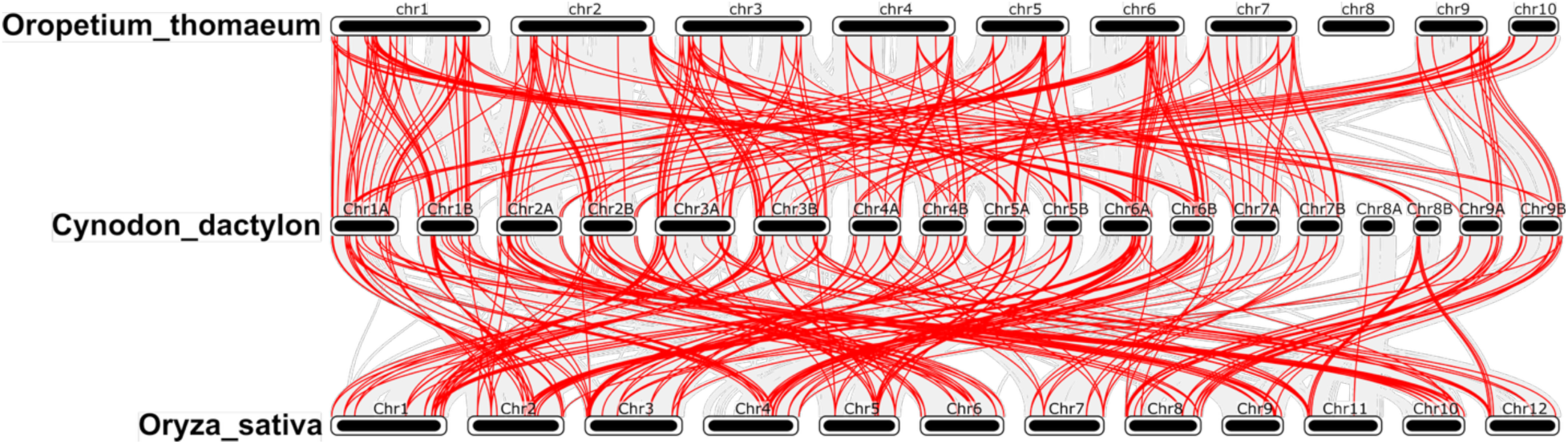
Syntenic analysis of CdNAC genes with *Oropetium thomaeum* and *Oryza sativa* genes

### Cis-element analysis of CdNACs

The cis-regulatory elements of all CdNAC genes were screened within a 2000bp upstream sequence of the start codon using PlantCare software. We identified 106 cis-regulatory elements (CRE) with known functions, which were broadly classified into 6 categories based on their major role (Table S3). 33 CRE were classified under light-responsive elements, including G-box, GT1-motif, and Box 4 TCT-motif. 17 CRE were grouped under hormone-responsive elements that include ABRE, CGTCA-motif, ERE, GARE-motif, and TGA-element. Similarly, 22 CREs were development-related, including CAT-box, RY-element, and O2-site. Another 22 elements were stress-related, including DRE, LTR, and MYB-binding sites. 10 CREs were classified under the core promoter class that includes elements like TATA-box and CAAT-box. And 2 elements, Box III and HD-zip3, were classified under the protein-binding category. The number and presence of CREs from each category greatly varied in the promoter regions of each CdNAC gene, indicating their differential regulatory potential and functional diversities (Figure S4). Excluding the core promoters like TATA-box and CAAT-box, 41% of the identified numbers were from stress-related class, 26% from hormone-responsive class, 19% from light-responsive class, and 9% from development-related class.

### Tissue-specific expression of the CdNAC gene family

The expression of CdNAC genes in different bermudagrass tissues, like inflorescence, root, leaf, rhizome, stolon, and shoot, was estimated using tissue-level expression data retrieved from the NCBI SRA database (PRJNA685207). Trimmed Mean of M-values (TMM) normalized counts per million (cpm) were used to study the relative CdNAC transcript abundance across different tissues in bermudagrass (Table S4). 105 CdNAC showed little or no detectable expression across the tissues tested, while the remaining genes showed moderate to high expression in at least one tissue. Among them, 57 CdNAC genes exhibited expression across all tissues, suggesting potential roles in fundamental biological processes. CdNAC6, 7, 12, 28, 50, 62, 77, 146, 153, 184, 222, 227, and 233 showed relatively higher expression in the root, possibly indicating root-related functional specialization (Figure 6). CdNAC5, 27, 96, 121, 182, 186, 187, 196, and 202 were predominantly expressed in the inflorescence and might have an important role in reproductive development. These patterns reflect the diverse functional and regulatory roles of NAC genes in bermudagrass.

**Figure 6:**
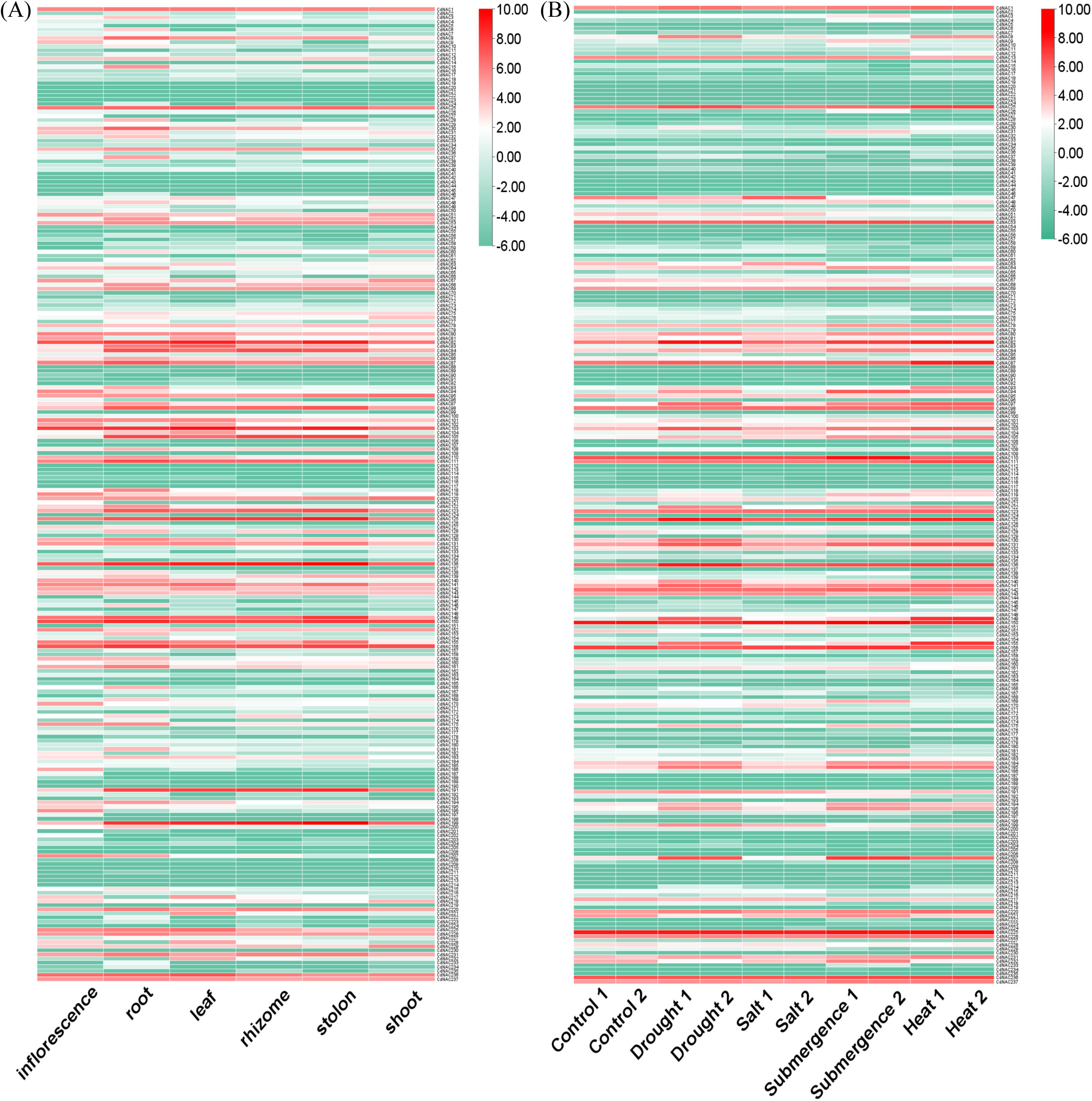
Heat maps showing log2 TMM normalized tissue level expression (A) and different stress level expression (B) of CdNAC genes

### CdNAC gene expression profile under different abiotic stresses

The role of CdNAC genes in abiotic stresses was evaluated by analyzing the differential expression of genes across drought, salt, heat, and submergence conditions as compared to control using RNA-Seq data (PRJNA589598). Differentially expressed genes (DEGs) were reported as significant when log_2_ fold change was >1 or <-1with a false discovery rate <0.05 (Table S5). The heat map in Figure 6 shows expression patterns under different stress conditions, while the Venn diagram in Figure 7 summarizes the DEGs. A total of 35, 43, 42, and 10 CdNAC genes were upregulated under drought, heat, submergence, and salt stress, respectively, and more than 50% of these genes were classified under the SNAC and NAM/NAC1 group in the above phylogenetic classification.

**Figure 7:**
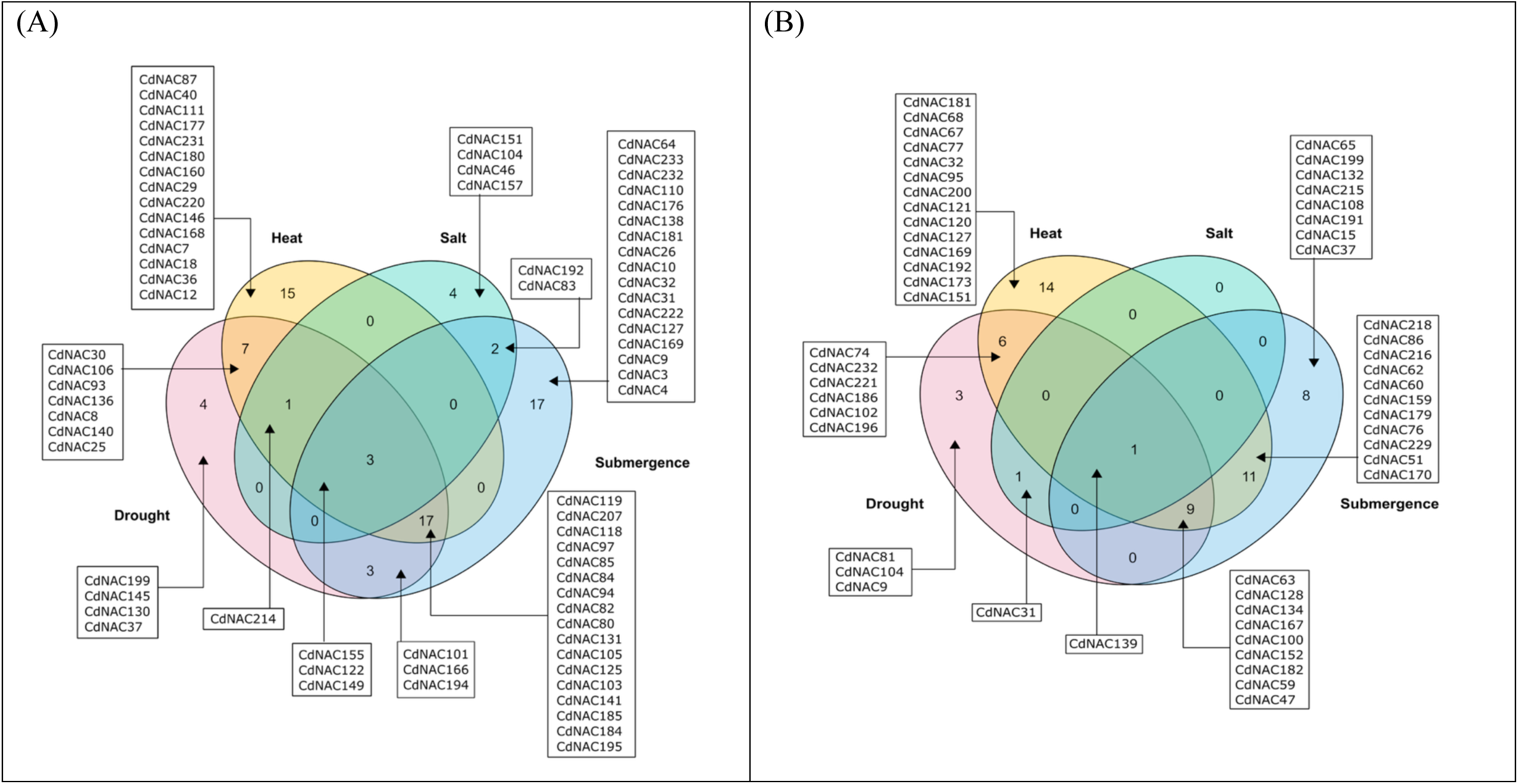
Venn diagram showing upregulated (A) and downregulated (B) genes across different abiotic stress conditions

CdNAC122, 149, and 155 were common to all stress conditions tested (Figure 7). CdNAC122 showed log2 fold changes (log2FC) of 4.37, 3.94, 1.2, and 3.04 under drought, heat, salt, and submergence stress conditions, respectively, compared with the control (Table S5). CdNAC149 showed log2FC of 5.86, 7.06, 2.03, and 3.09 during drought, heat, salt, and submergence stress compared with control conditions. CdNAC155 had a log2FC of 6.36, 7.94, 2.45, and 3.93 during drought, heat, salt, and submergence stress compared with the control. While CdNAC37, 130, 145, and 199 were drought specific, CdNAC46, 104, 151, and 157 were salt specific, 15 genes, including CdNAC7, 12, 18, and 29, were heat specific, and 17 genes, including CdNAC3, 4, 10, and 26, were submergence specific (Figure 7). The XND1 group genes, like CdNAC9 and CdNAC31, had unique expression in submergence stress, potentially supporting its root hydraulics and stress trade-off functions. Similarly, 20, 41, 29, and 2 genes were downregulated under drought, heat, submergence, and salt stress, respectively. Most of these genes belong to classes NAM/NAC1, ONAC022/TERN, and OsNAC7, well known for their meristem activity, organogenesis, stress-responsive regulators, and developmental processes.

## Discussion

Common bermudagrass is one of the few plant species with worldwide distribution. Its cosmopolitan nature is attributed to its high resilience to environmental stresses [8,9]. Its ability to thrive in high temperatures, survivability in marginal land with low water requirement, ability to tolerate moderate salinity, and endurance to short periods of flooding make it a desirable grass for soil stabilization, turf, and forage purposes. These adaptive characteristics suggest the involvement of numerous genes contributing to the stress resilience of this species.

Since the first discovery of NAM family genes in petunia, their study has gained popularity due to their critical role in plant growth and development, including shoot apical meristem and primordial boundaries formation [28]. Subsequent discovery of stress-responsive ATAF1/2 genes and development-related CUC2 genes comprising a similar structure, including conserved N-terminal sub-domains, gave rise to the NAC gene family: one of the largest transcription factor gene families in the plant kingdom [29]. Since then, a comprehensive analysis of this gene family has been done in many plant species for their functional studies.

In this study, we identified 237 NAC transcription factors in common bermudagrass. The large number of NAC genes in bermudagrass may be attributed to polyploidy nature of the species, which is in accordance with a high number in other polyploid species, like 251 NAC genes in switchgrass, 333 in oat, as compared to diploid species, like 140 in rice, 117 in *Arabidopsis* [25,27,30,31]. Based on sequence similarity, we have classified 237 CdNAC genes into 14 classes, including identification of the 23 members belonging to SNAC (AtNAC3/ATAF/NAP), which have an important role in abiotic stress tolerance. The large number of collinear gene pairs, both within bermudagrass and close grass relatives like resurrection plant and rice, suggests whole genome or segmental duplication as the primary mechanism of gene family expansion. Cis regulatory element analysis from the promoter region of the genes returned 41% (excluding core promoter elements) cis elements from the stress-responsive class, indicating a critical role of CdNACs in stress responses.

Expression profiling revealed that more than 50% of the CdNAC genes were expressed in at least one of the tissues assessed. 9 inflorescence-specific CdNAC genes also highlight the importance of CdNAC in flowering and fruiting. J. Liu et al. (2023) has reported regulatory roles of NAC genes in the expression of the inflorescence formation-related genes, potentially through controlling the transition of the shoot apical meristem to the floral meristem. Tang et al. (2023) reported overexpression of ZaNAC93 in tomato induced early and increased flowering, through GA, ABA, and JA signaling pathways that contribute to floral induction. Similarly, expression of 13 CdNAC genes only in roots also highlights the important role of NAC genes in root architecture and development. Hao et al. (2011) found overexpression of GmNAC20 genes promoting lateral root formation by altering auxin signaling-related genes in transgenic soybean. P. Xu et al. (2022) identified nitrate-inducible NAC056 promoted nitrate assimilation and root growth in *Arabidopsis*.

Similarly, we identified 75 genes upregulated and 53 genes downregulated under at least one of the drought, heat, salt, or submergence stress conditions. Several studies have supported the crucial role of NAC genes in abiotic stress responses, and their overexpression has resulted in improved abiotic stress tolerance in plant species. For example, overexpression of ANAC019, ANAC055, ANAC072, and ATF1 in *Arabidopsis* [36,37], OsNAC6, ONAC045, ONAC022, and ONAC066 in rice [38–41], and ZmNAC20 in maize [42] resulted in improved drought and salinity tolerance through mechanisms such as osmotic adjustment and ABA signaling. Overexpression of ONAC003 in rice (Y. Fang et al., 2015) ATAF1 and ANAC055 in *Arabidopsis* [44,45] conferred heat stress tolerance through ROS scavenging and heat shock pathways, while ANAC102 in *Arabidopsis* mediated flood stress through ABA and ethylene signaling pathways [46]. These findings suggest that the broad expression of NAC genes might have contributed to the resilient nature of bermudagrass, and their overexpression could provide new opportunities for enhancing stress tolerance in bermuda as well as other grass species. However, the mechanisms and pathways of both upregulation and downregulation of NAC genes during different abiotic stress conditions require further comprehensive functional studies.

## Conclusion

This study provides the first comprehensive genome-wide analysis of the NAC transcription factor gene family in common bermudagrass, identifying 237 CdNAC genes. The expansion of this gene family is likely linked to the gene duplication and polyploid nature of the species and reflects the complex evolutionary landscape of its genome. Phylogenetic classification, cis-regulatory elements analysis, and tissue-specific expression profiling revealed that CdNACs are involved in a wide range of biological functions. Differential expression of CdNAC genes over the period of abiotic stress, like drought, salt, heat, and submergence, identified CdNAC genes particularly involved in abiotic stress response. The findings may lay the foundational work for future functional characterization and verification of NAC genes and their application in breeding of abiotic stress resilient bermudagrass.

## Data Availability Statement

All datasets analyzed in this study are publicly available. Bermudagrass genome sequence and annotation were obtained from Genome Warehouse under the assembly number PRJCA019991. Raw expression data were retrieved from the NCBI SRA database under the accession numbers PRJNA685207 and PRJNA589598. This study presents the first genome-wide identification and characterization of CdNAC genes, and the curated gene dataset and expression profiles are provided in Supplementary tables.

## Author Contributions

AP: Conceptualization, Data curation, Methodology, Formal analysis, Writing – original draft. YW: Conceptualization, Methodology, Resources, Supervision, Writing – review & editing.

## Acknowledgements

We would like to acknowledge Dr. John E. Gustafson of the Department of Biochemistry and Molecular Biology, Oklahoma State University, for his support and guidance throughout this study. We also appreciate technical support from the High-Performance Computing Center (HPCC) at Oklahoma State University.

## Supplementary Information

Supplementary_Figures.docx: Figure S1 – S4. Supplementary_Tables.xlsx: Table S1 – S5.

