## Supplemental Data 1 for "Genome-wide identification and characterization of the NAC transcription factor family in *Cynodon dactylon* and their expression during abiotic stresses"

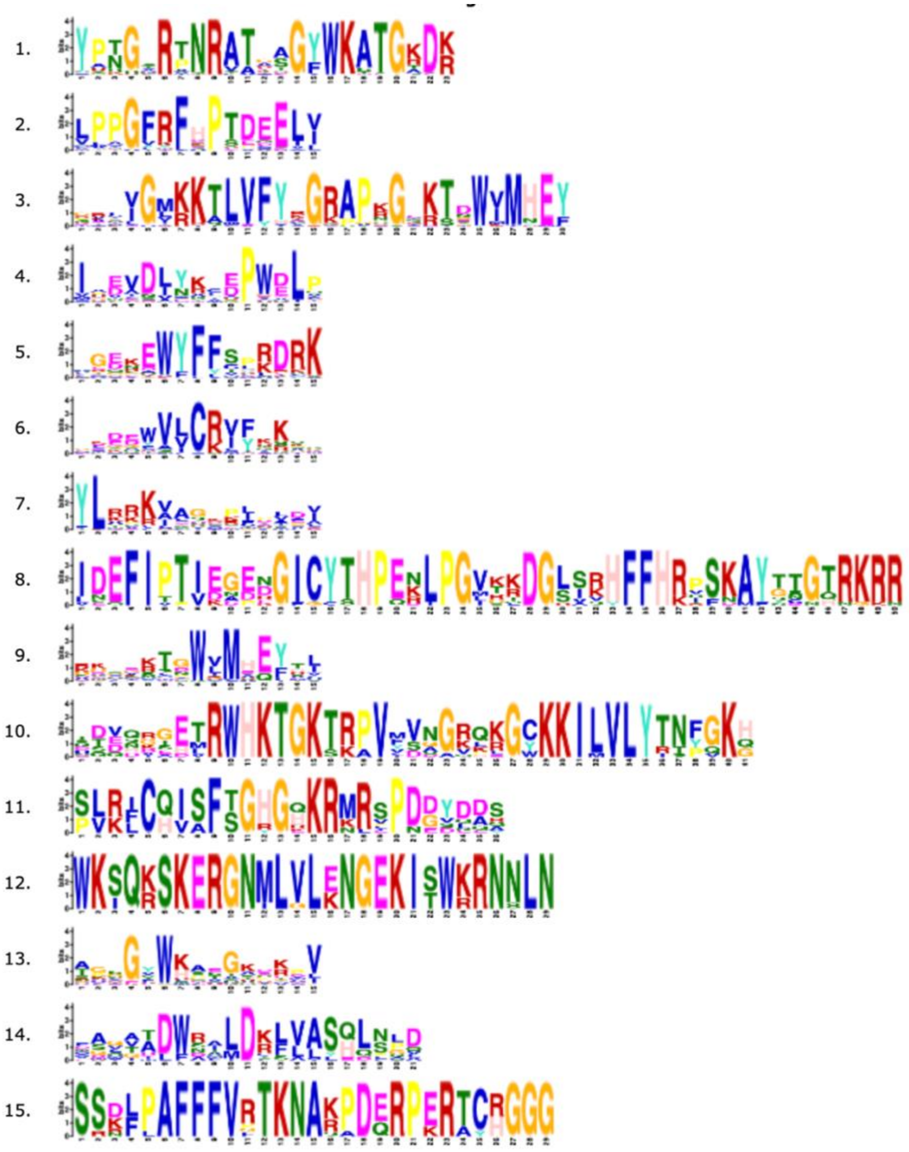

Figure S1: Sequence logos of the top 15 motifs in CdNAC proteins discovered using MEME

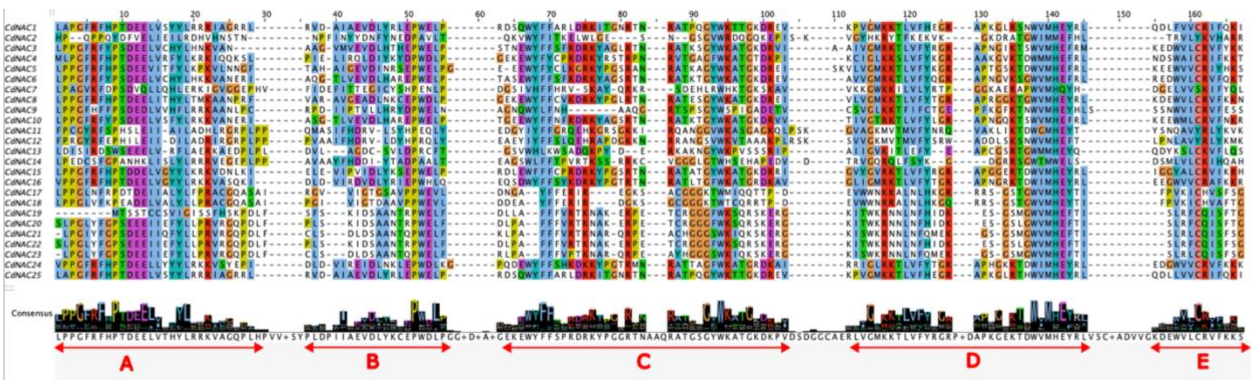


Figure S2: The representative five N-terminal sub-domains of the CdNAC proteins

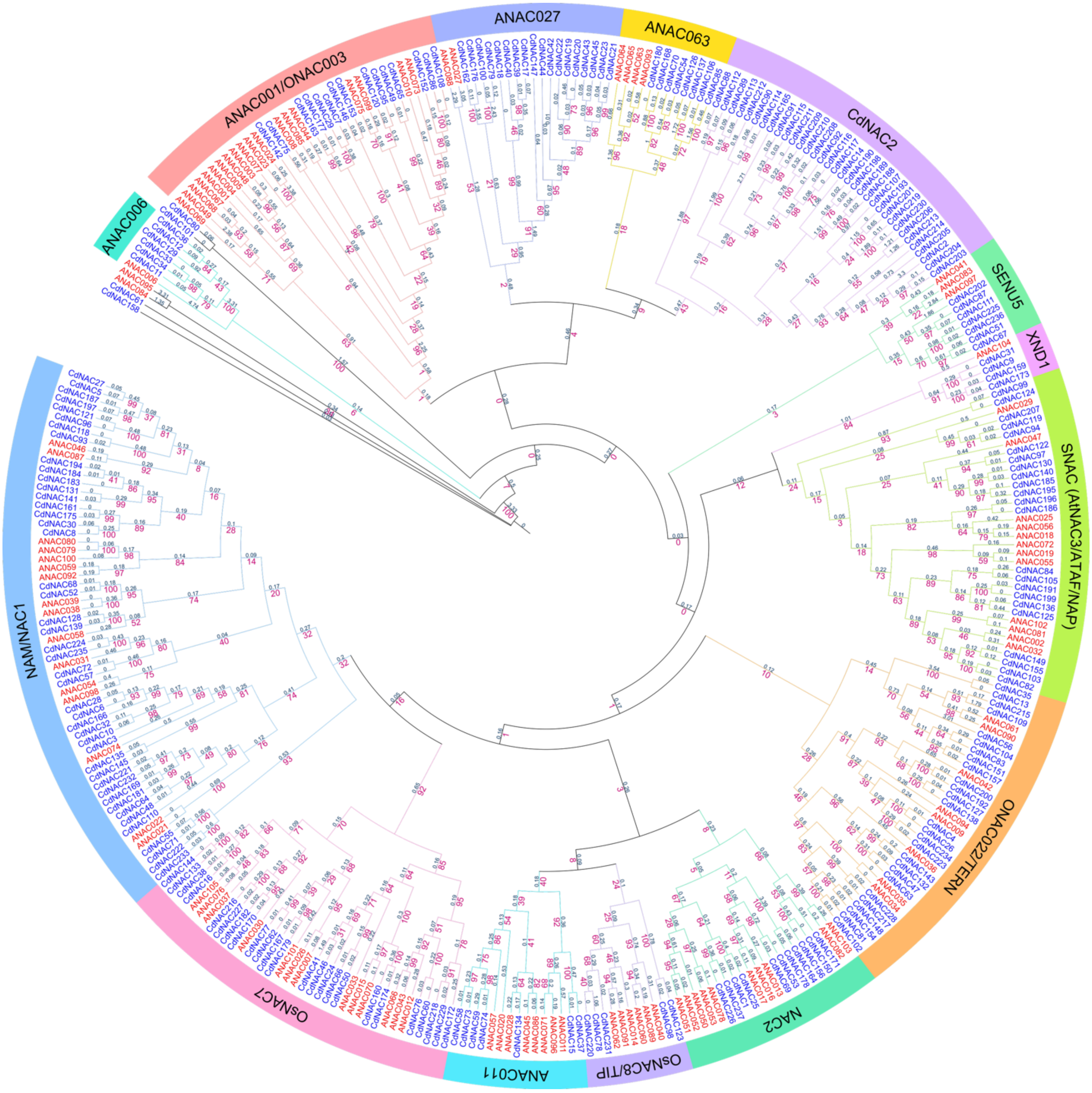


Figure S3: Phylogenetic tree of bermudagrass and *Arabidopsis* NAC genes with bootstrap support values and branch length

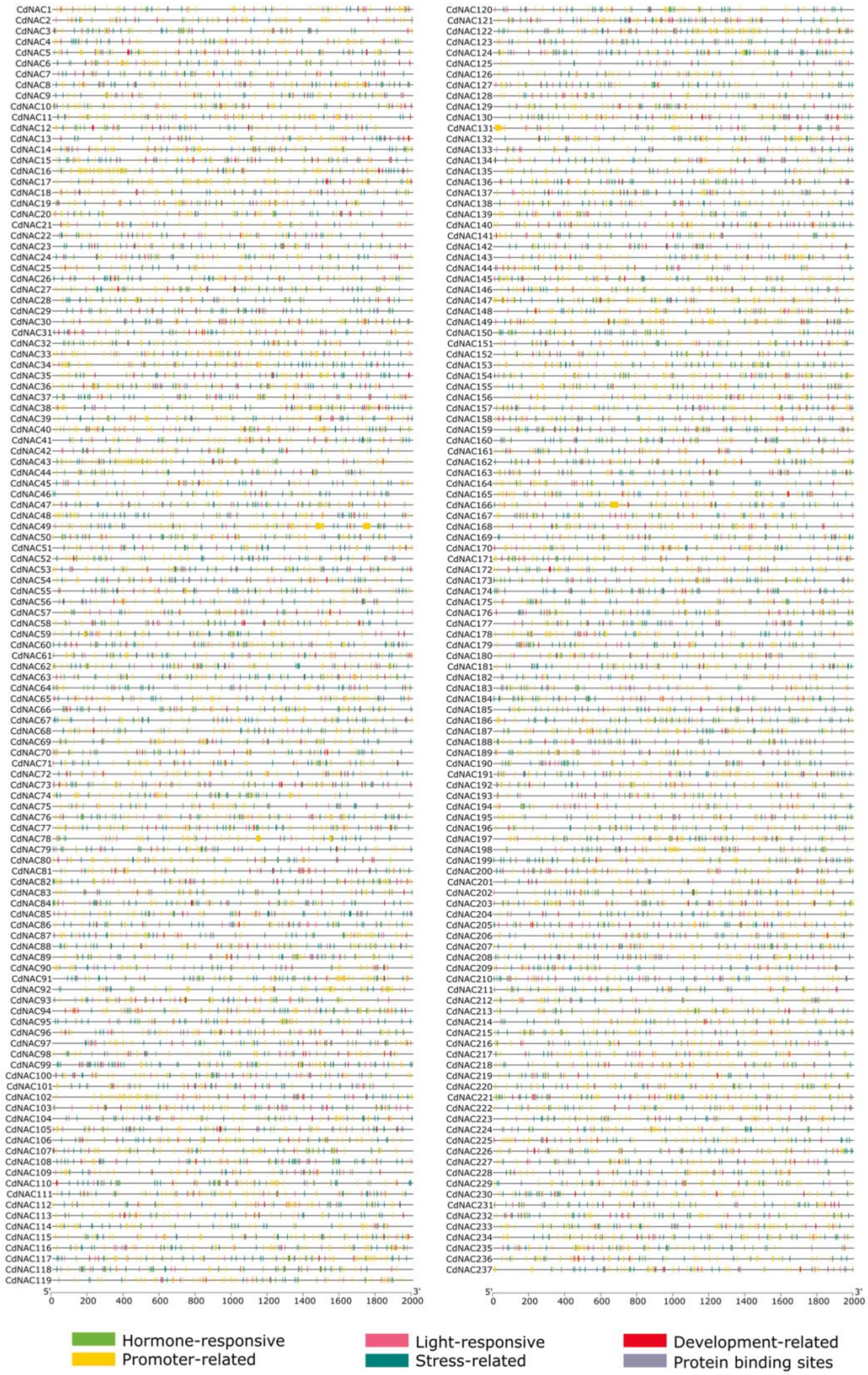


Figure S4: Cis-regulatory elements in the 2kb upstream region of CdNAC genes
